# Correlation between bioluminescent blinks and swimming behavior in the splitfin flashlight fish *Anomalops katoptron*

**DOI:** 10.1101/2024.02.21.581075

**Authors:** Peter Jägers, Timo Frischmuth, Stefan Herlitze

## Abstract

The light organs of the splitfin flashlight fish *Anomalops katoptron* are necessary for schooling behavior, to determine nearest neighbor distance, and to feed on zooplankton under dim light conditions. Each behavior is coupled to context-dependent blink frequencies and can be regulated via mechanical occlusion of light organs. During shoaling in the laboratory individuals show moderate blink frequencies around 100 blinks per minute. In this study, we correlated bioluminescent blinks with the spatio-temporal dynamics of swimming profiles in three dimensions, using a stereoscopic, infrared camera system. Groups of flashlight fish showed intermediate levels of polarization and distances to the group centroid. Individuals showed higher swimming speeds and curved swimming profiles during light organ occlusion. The largest changes in swimming direction occurred when darkening the light organs. Before *A. katoptron* exposed light organs again, they adapted a nearly straight movement direction. Light organs create a strong contrast against the background. Therefore, a close combination of light signals and movement is crucial to the behavior of *A. katoptron*.

## Introduction

Theoretical [1] and experimental [2] observations of animal movement range from individual levels of interaction (e.g. vortex phase matching [3]) to large-scale phenomena (e.g. migration [4]). In fish, motion is ubiquitous and depends on numerous external (e.g. predatory pressure or environmental stress; [5]) and internal (e.g. genetic or physiological; [6]) factors. To enhance fitness in a changing environment or under threat, motion is necessary. This becomes obvious in the context-dependent movement profiles such as startle responses [7], freezing [8] or unpredictable, erratic changes of swimming direction that can help to distract predators ([9, 10]; also see [11]).

Most animals adjust their speed and turning rates to regulate their movement direction. The forces of motion acting on the bodies can be determined with mathematical approaches [12]. For example, fish show decreased turning rates at higher swimming speeds due to inertial restrictions [13]. Maximum swimming speed of marine fish depends on tail beat frequency, body length and hydrodynamic environment [14] and can exceed 10 body lengths/s [15]. Swimming speed is limited to increased oxygen consumption and muscle tonus, and needs to be balanced against putative advantages such as finding new resources [16]. The lateral line is essential for monitoring the hydrodynamic properties of the environment and it plays a crucial role in sensing group members while schooling [17].

Living in groups provides numerous benefits to fish (e.g. vigilance, reproductive success or energetic benefits) [18], but requires increased group coordination to avoid collisions. The advantages frequently rely on the ability of group members to share information, allowing for collective decisions and maintaining group coordination. Whether under predation risk or during foraging, information transfer based on intentional signals or passive cues is an essential element [19]. In fish, information is detected using a range of sensory modalities, with the focus in previous studies having been on vision [20], with far fewer studies on sound [21], olfaction [22], and electrocommunication [23].

The spatial and temporal organization of fish shoals shows a strong variability from highly polarized to dispersed motion [24, 25]. Polarized movements are frequently observed during fast escape responses and reduce the risk of predator attacks [26]. In contrast, slow moving groups are more dispersed, thereby, increasing visual fields with higher probabilities to spot threats or resources. The transition from ordered to disordered motion is dependent on context [27], moving speed [28, 29] or group densities [30]. Other effects like lateralization can also be associated to collectively moving fish groups [31].

More recently, different technologies (full spherical imaging [32]; or sonar [33]) were established to observe large scale movement profiles. In our study, we used a stereoscopic, infrared camera system to record movement profiles and bioluminescent blinks of the group-living flashlight fish *Anomalops katoptron*.

Bioluminescence shows a high abundance in the ocean and has multiple functions e.g. to conceal the body contour via large amounts of photophores [34] or to create point-like light sources in visually restricted environments [35]. The nocturnal flashlight fish *A. katoptron* (Anomalopidae) inhabit the Indo Pacific and appear in aggregations ranging from small, uncoordinated to large, polarized groups [36]. Characteristic for the Anomalopidae is a sub-ocular light organ, which is densely packed with bioluminescent, symbiotic bacteria [37]. *A. katoptron* exhibit a downward rotation of the light-emitting surface to shield the bacteria’s continuous illumination [38]. By alternating light organ occlusion and exposure, individuals create distinct, context-dependent blink patterns, which have been shown to be involved in the localization of zooplankton [36], orientation towards conspecifics [39], and intraspecific communication [40]. Most bioluminescent fish species show adaptations in the visual systems e.g. intraocular filters [41] or possess opsins optimized to visualize the luminescent signals [42]. Opsins of flashlight fish show the highest sensitivity within the frequency range of ambient star/moon-light and the emission spectra of their own symbionts (480-510 nm) [43]. They are not able to recognize longer wavelengths such as infrared light and, therefore, can be monitored with infrared camera setups.

Besides offensive functions of bioluminescence (e.g., prey attraction), defensive functions such as screens, predator distraction, counterillumination or startle have been described [35, 44]. However, description of movement profiles in combination with bioluminescent flashes of nocturnal, marine organisms remain scarce. In fish, a distraction of predators has been proposed for *Gazza minuta* [45] and *Photoblepharon steinitzi* (Anomalopidae) [46]. Furthermore, bioluminescent backlighting in the Humboldt squid *Dosidicus gigas* is combined with specific locomotor behaviors and has been proposed to facilitate intraspecific communication. Additionally, these behavioral patterns, although filmed during the day, have been proposed to distract predators while being temporarily vulnerable during hunting [47]. The pattern of precisely timed bioluminescent flashes and movement profiles of the male ostracod *Photeros* (formerly *Vargula*) *annecohenae* has been linked to sexual courtship [48].

Our study focusses on individual movement dynamics in three dimensions in a visually restricted environment. We show that *A. katoptron* rapidly changes swimming direction in correlation with the blink patterns of its light organs. In addition, individuals show increased swimming speeds with occluded light organs.

## Results

To investigate the correlation of the light organ occlusion/exposure with the movement profiles of the nocturnal flashlight fish *Anomalops katoptron*, we recorded three dimensional trajectories of small groups of *A. katoptron* under infrared settings (Fig. 1 and Additional File 3). To determine attributes of shoaling, we analyzed polarization, the alignment of individuals, and mean distance to the group’s centroid. Individuals within a group of *A. katoptron* did neither move in the same (values 1) nor opposite (values 0) direction (Fig. 2).

**Figure 1:**
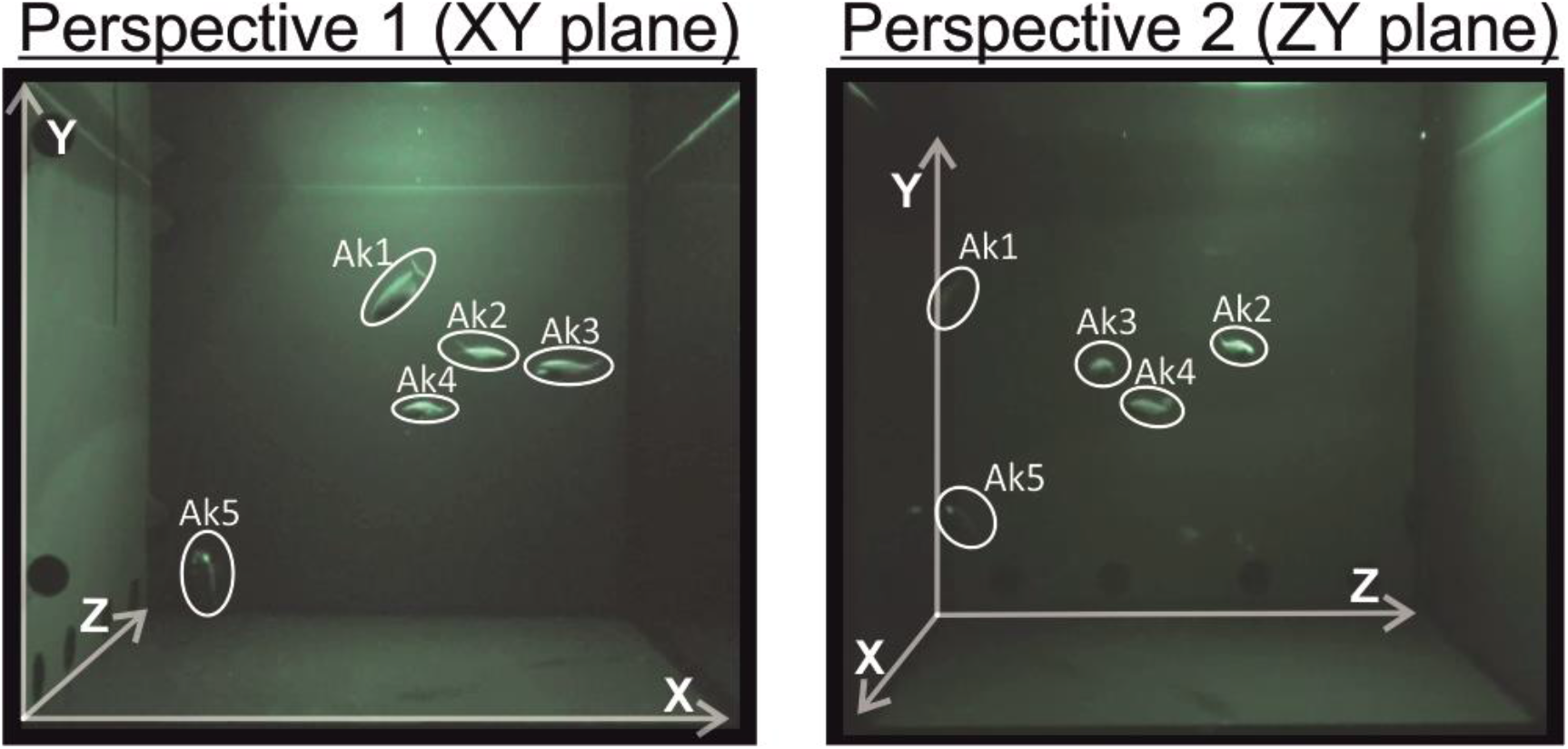
Camera Perspectives. To achieve three-dimensional tracking profiles, we used two infrared camcorder arranged in orthogonal orientation. Shown are both (XY- and ZY) planes.

**Figure 2:**
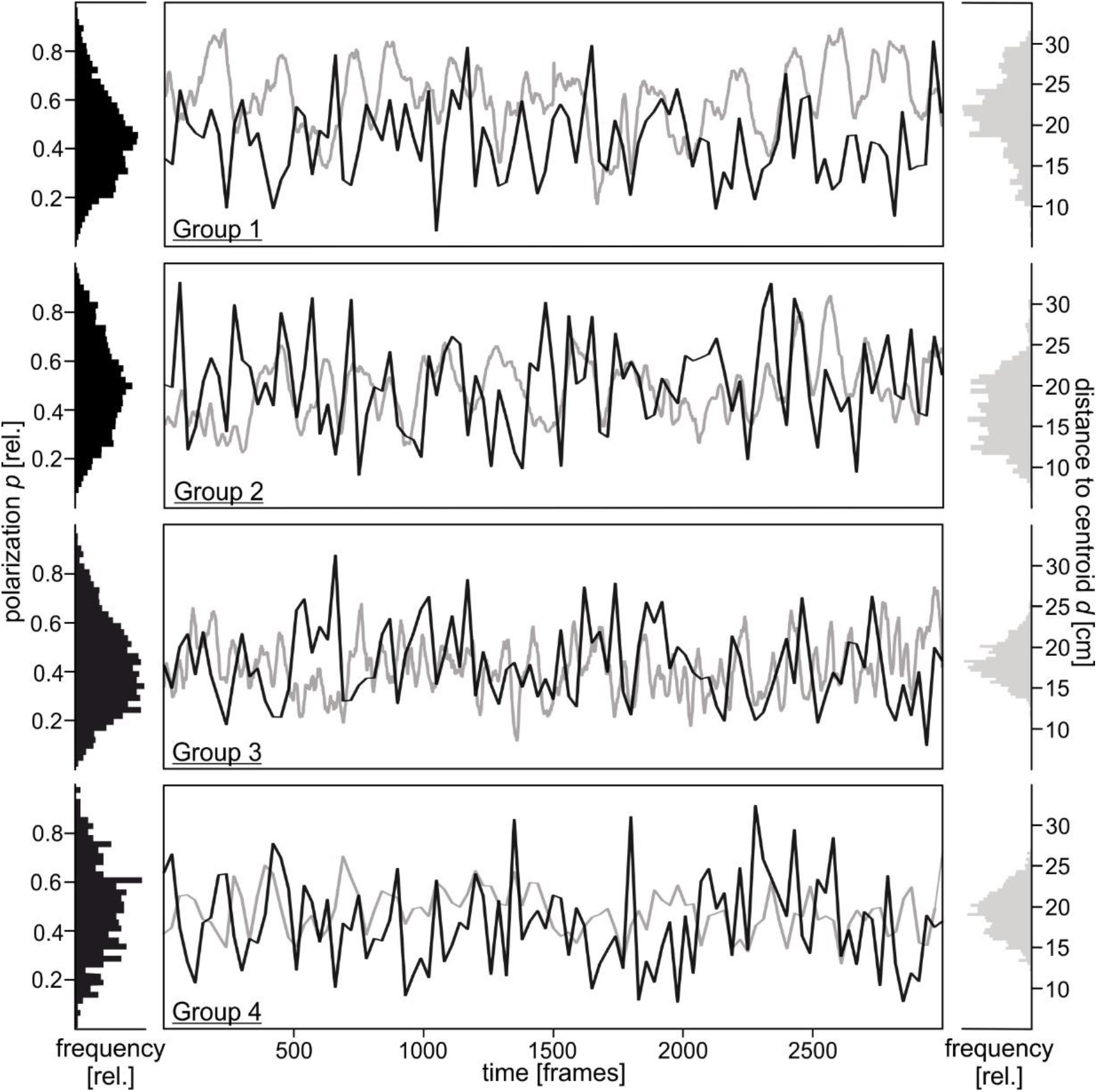
Shoaling of *A. katoptron*. Trajectories of four groups, each consisting of five individuals, were obtained from two camera perspectives. Polarization (*p*, black) and distance to the group’s centroid (*d*, grey) was characterized for 3000 frames (B). Histograms (bin size 40) show densities of polarization and distance to centroid for the full recording time (3000 frames). Polarization was smoothed via averaging the values at neighboring points (sampling proportion 0.001 ≙3 frames; fraction of a total number of data points used to compute each smoothed value).

Additionally, we discovered that the distance of *A. katoptron* to the centroid occurred in a wave-like pattern and the mean polarization was moderate (Fig. 2). In our study, the mean light organ exposure (345 ± 14.7 ms) is different from the occlusion (245 ± 18.3 ms; *t*(19) = 3.489, *p* = 0.002; Fig. 3B). The alternating exposure and occlusion resulted in blink frequencies of 103.82 ± 4.15 blinks/min.

**Figure 3:**
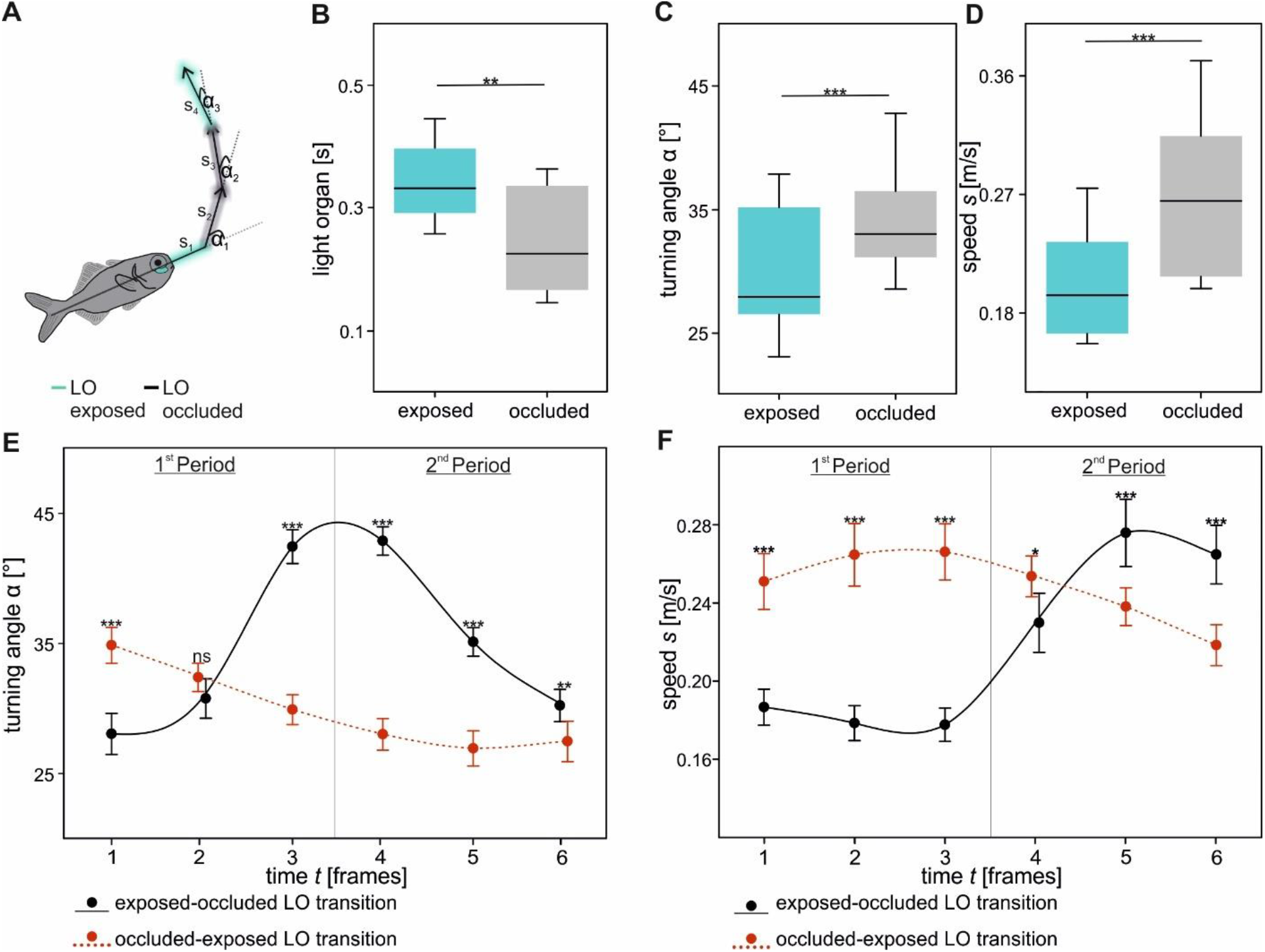
Individual movement direction in relation to light organ (LO) exposure and occlusion. Example trajectory of one individual (A) and light organ exposure and occlusion during shoaling (B). Differences of swimming direction (C) and speed (D) under both conditions (LO exposed or occluded). Detailed changes of swimming direction (E) and speed (F) three frames before (1-3) and after (4-6) the light organ transition. Data was obtained from twenty individuals (*n =* 20). A dynamic fitting with a polynomial, cubic equation was used to additionally plot data in E and F. Significance values are reported as *p < 0.05, **p < 0.01, ***p < 0.001. Data in E and F indicate mean ± SEM.

Individuals showed larger turning angles (34.31 ± 1.04 °; *t*(19) = - 7.94, *p* ≤ 0.001; Fig. 3C) and increased swimming speeds (0.267 ± 0.014 m/s; Wilcoxon signed rank: Z = 3.92, *p* ≤ 0.001; Fig. 3D) with occluded compared to exposed light organs. A more detailed analysis revealed the differences in swimming direction and speed between the transition from exposed to occluded light organs and vice versa (Fig. 3E and F). The strongest alteration became obvious when light organs were darkened. Immediately before occluding their light organs, individuals slowed down to 0.178 ± 0.008 m/s. This was combined with a change in swimming direction around the transition (42.447 ± 1.3 °, frame 3 and 42.879 ± 1.1 °, frame 4). In the consecutive frames, individuals increased swimming speed (0.265 ± 0.015 m/s, frame 6) and decreased swimming angle to 30.225 ± 1.24 ° (frame 6).

The transition from occluded to exposed light organs showed smaller alterations, indicating a continuous, straight-lined swimming profile. The swimming speed was slightly increased to approx. 0.25 m/s during the transition (frame three and four). Turning angle was nearly constant with sustained light organ exposure in frame five (26.93 ± 1.347 °) and six (27.469 ± 1.553 °).

Frame related transitions of occluded to exposed and exposed to occluded light organ were significantly different regarding turning angle (Fig. 3E; F 5, 95 = 63.61, p ≤ 0.001; Table S1) and swimming speed (Fig. 3F; F 5, 95 = 38.72, p ≤ 0.001; Table S2). The post-hoc analysis revealed that besides frame two for turning angle (Holm-Sidak: p = 0.117), all other results were highly significant (Holm-Sidak: p ≤ 0.001).

In summary, flashlight fish *A. katoptron* coordinate bioluminescent blinks with changes in movement profiles while shoaling.

## Discussion

In the present study, we observed that *Anomalops katoptron’*s blink and movement profiles followed a precisely timed pattern while shoaling. When individuals occluded their light organs, their swimming speed increased, and swimming direction was changed.

Shoals of *A. katoptron* can be observed during moonless nights at the water surface of the Indo Pacific [49]. During shoaling, each individual uses sub-ocular light organs to emit bioluminescent blinks [40]. We found that small groups of *A. katoptron* showed mean blink frequencies of 103 blinks/min with slightly increased light organ exposure compared to occlusion. Occasionally, we observed that light organs of an individual were not exposed simultaneously.

Bioluminescent signaling can enhance intraspecific communication in visually restricted environments as shown in ostracods (e.g. Cypridinidae) [48], cephalopods (e.g. Ommastrephidae) [47], and fish (e.g. Leiognathidae) [45]. For group living *A. katoptron*, bioluminescent displays have been proposed to attract conspecifics [39], determine nearest neighbor distance [40], and illuminate prey [36]. Other than *A. katoptron*, individuals of the closely related genus *Photoblepharon* occur solitary or in pairs. Here, the distraction of predators via “blink and run”-pattern [46], aggression during territorial defense, and illumination of prey [50] have been linked to bioluminescent displays. Besides the benefits of bioluminescent signaling, light sources also build a strong contrast against a dark background (e.g. with increasing water depth), becoming increasingly visible to predators with appropriate visual systems [51]. To balance the visual information for conspecifics while reducing the risk of being exploited by predators is crucial.

Our study revealed that after light organs were occluded, individuals immediately changed direction and increased swimming speed. This provides coverage for the last visual cue’s spatial position.

The confusion of predators has been proposed for bioluminescent fish species such as *Gazza minuta* [45], *Leiognathus elongatus* [52] (note taxonomic revision [53]), and the “blink and run” pattern of the related *Photoblepharon steinitzi* [46]. Our results agree with the “blink and run”-hypothesis, where evasive swimming and bioluminescent blinks are coordinated [46]. For other, non-bioluminescent fish species it has been suggested that a rapid change of movement direction increases survival during predator attacks. For example, virtual prey with straight swimming trajectories (Lévy motion) were targeted more frequently by predators [54]. In addition, several other defensive functions of bioluminescence e.g. burglar alarm, distractive body parts or smoke screens have been discussed in many other species [35].

Besides the suggestion of a distractive signal, visual cues can also be important for group coordination. It has been shown that bioluminescent signals of *A. katoptron* are necessary to school under dim light conditions and small numbers of individuals can initiate changes in movement directions [39]. While schooling, for example during fast escape responses, individuals are synchronized in their movement and blinking pattern [39, 40]. Swimming speed has been positively correlated with higher group polarization in other species [55]. Furthermore, because of inertial limitations, turning rates are reduced at faster swimming speeds [13]. In contrast to coordinated group behavior, our results show low polarization and swimming speed of groups of *A. katoptron* while shoaling in the tank.

Limitations due to the spatial constraints of the tank walls which may affect the acceleration and turning response of the individuals are possible. Other species maintain a minimum distance of 5 cm towards the wall [2]. Conversely, we observed that *A. katoptron* sometimes moved in front of the mirrored tank wall in response to their own reflection (see Additional File 3). The perception of an attracting signal is most likely (similar to [40]; also note self-recognition in other species [56]).

Our study is limited to small groups of *A. katoptron* under laboratory conditions. Furthermore, our manuscript does not explore the trade-off between distraction of predators and attraction of conspecifics. This needs to be addressed in future work, focusing on the information transfer during the transition from loosely organized to highly synchronized schools. Here, tracking software based on high-resolution, deep-learning approaches [57] and advanced technological approaches within field sites would be necessary [32]. Information transfer during transitions have been observed either in non-bioluminescent fish species e.g. *Notemigonus crysoleucas* [58] or terrestrial environments e.g. fireflies *Photinus carolinus* [59].

In summary, our results show that individuals of *A. katoptron* correlate directional changes and light organ occlusion during shoaling. This may suggest that bioluminescence in *A. katoptron* is important not only for intraspecific communication [40], but also might serve as a defensive mechanism.

## Methods

### Husbandry

Specimen of *Anomalops katoptron* (n = 20; total body length: 7.71 ± 0.08 cm) were obtained from DeJong Marinelife (Netherlands). Animals were maintained for several weeks before the experiments were carried out.

In the laboratory, the light-dark cycle was set to 12 h – 12 h with the dark period starting at 12 h am CET. During the day, groups of *A. katoptron* dwell in caves and crevices with low light intensities [40]. Therefore, we placed different shelter in the tank and installed opaque PVC cover around it. The housing tank (120 cm x 60 cm x 60 cm; L x W x H) was equally subdivided by an opaque PVC plate. The compartments (58 cm x 58 cm x 55 cm; L x W x H) were connected via a sliding door and individuals were allowed to switch between sides. Standardized filter systems and aeration were used (see [36, 40] for details) to achieve steady water parameters (temperature: 25-27 °C; salinity: 34-36 ‰; NO3 < 20 mg/l; NO_2_ < 0.1 mg/l; PO_4_ < 0.1 mg/l). Once a day, short periods (< 30 seconds) of dim red light (TX 100; Coast; USA) were used to illuminate the tank and individual health was observed. Twice a day, individuals were fed ad-libitum under dark conditions with defrosted zooplankton and small amounts of minced salmon.

### Experimental procedure

For the experiments, one compartment of the tank was emptied, and the opaque sliding door was closed. The tank was illuminated with overhead infrared torches (ʎ_max_= 850 nm; IKV ACC-07, Inkovideo GmBH, Germany). Other light emitting sources were turned off or darkened. The experiments began at 2 pm (CET), two hours after the dark period started.

To achieve a stereoscopic view, two, infrared-sensitive camcorder (HDR-CX730, Sony, Japan) filming with a resolution of 1920 × 1280 pixel at 25 fps were placed in an orthogonal orientation in front and at the side of the compartment (Fig. 1 and S1A). Each camcorder was placed on a tripod at the same level (160 cm in our setup) and 77 cm from the tank, matching the center of the compartment. We used the Stereo Camera Calibration toolbox of Matlab (Matlab 2022b; The MathWorks Inc., USA) to compute camera parameter by taking various photographs (n = 16) of a checkerboard (7 × 9; squares: 30 × 30 mm; Fig. S1A) in different orientations. The mean projection error was 0.71 pixel (Fig. S1B). The toolbox allowed us to determine intrinsic parameters for both cameras, as well as the rotation and translation of camera two in reference to camera one, which was designated as the scene’s center. Further, we calculated projection matrices for both cameras (Additional File 1; Fig. S1C).

Prior to the experiment, visibility of the tank was checked through the ocular, which then was darkened with black sheets. For the experiment five individuals were randomly assigned to a group. The first group was measured on the 21st June 2021, the second group on the 31st August 2021, and groups three and four on the 7th September 2022. Each group was transferred with a small hand net to the measurement compartment and habituated for five minutes in total darkness. Camcorder were started and recordings took place for five minutes. After recording, individuals were transferred back into the housing compartment.

### Data analysis and Statistics

We back-synchronized camcorder with brief (< 1 s), dim, red light pulses presented after each trial. Additionally, we checked multiple frames in which light organs were visible in both perspectives and controlled whether synchronization occurred. Videos were converted to .avi-format and edited by using Shotcut (GNU General Public License; Meltytech, LLC). Two minutes (≙3000 frames) of swim and blink profiles of twenty *A. katoptron* were manually analyzed (total of 4163 blink events), frame by frame in Vidana (Vidana 2.0, Germany). The x- and y-pixel coordinates were obtained from both cameras (Fig. 1). We used a script originally written by Lourakis [60] to triangulate real world coordinates (given in mm). Here, we used the midpoint method [61] to generate projection matrices (Fig. S1). The real-world coordinates were subsequently assembled to respective time frames in Excel (Office Professional Plus 2019; Microsoft, USA). Light organ exposure (assigned to 1) and occlusion (assigned to 0) were documented and added to the excel files as soon as one light organ was exposed (see Additional File 2).

To explain group features, we examined individual distances to the group’s centroid and polarization, which offers a measure of alignment of individuals inside the group. The value of the individual (*n*) heading at time point *t*, where *u*_*i*_ is the unit vector of fish number *i*, was used to derive polarization (*p*, Eqn 1).

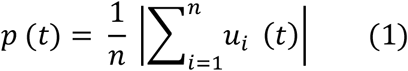

Values reached *p* = 1 when all individuals were aligned, whereas *p* = 0 when no alignment existed. The group’s centroid was determined as the mean of all individual coordinates of the group at timestep *t*. Respectively, distance to the group’s centroid (*d*) was calculated and averaged for all individuals within the group. Speed and change of swimming direction were calculated with self-written Matlab programs (Matlab 2020a; The MathWorks, Inc., USA). Thereby, instantaneous speed *s*(*t*) was the distances between coordinates at time *t* (Eqn 2).

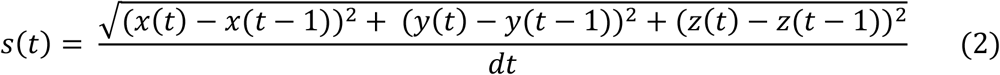

Here, *dt* is the length of the time interval (*dt* = 1/fps ≙0.04 s) and *x(t), y(t)* and *z(t)* are the *x, y* and *z* coordinates of one fish at time *t*. The change in direction (α) was calculated by the arccosine of two vectors (Eqn 3). Each vector (*u, v*) was determined for a pair of coordinates, vector *u* at timepoints (t), (t - 1) and *v* at timepoints (t), (t + 1). A 180-degree angle is equal to a U-turn, whereas 0-degree represents a straight line.

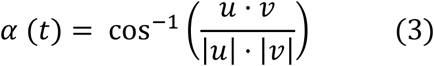

We analyzed speed and angular changes three frames before and after phase transition of light organ exposure to occlusion and vice versa. For every individual, we calculated the mean values at each time step. In Figure 3 E and F a dynamic fitting with a polynomial, cubic equation was used to plot data (included function of SigmaPlot; version 12.0; Systat, India).

The descriptive statistic (e.g. mean and standard deviation) was calculated in Excel. The data points of all individuals were pooled and analyzed in SigmaPlot. After successful evaluation of normal distribution (Shapiro-Wilk test), differences in exposure and occlusion of light organs (Fig. 3B), and angular changes (Fig. 3C) were analyzed with a paired t-test. In case of non-normally distributed data (swimming speed; Fig. 3D), the Wilcoxon signed rank test was used to assess differences. Differences of turning angles (Fig. 3E) and swimming speeds (Fig. 3F) at the specific timesteps were evaluated via a two-way repeated measurement ANOVA and Holm-Sidak post hoc analysis. The analysis was performed using timestep and type of transition (either exposed to occluded light organs or vice versa) as factors (see Table S1 and S2 for detailed values). All values are reported as mean ± SEM (standard error of mean). Significant differences are reported as: * p ≤ 0.05, ** p ≤ 0.01; *** p ≤ 0.001.

### Figures

Figures were created with SigmaPlot 12.0 (www.sigmaplot.co.uk) and Matlab (Matlab 2022b; The MathWorks Inc., USA), and processed with CorelDraw Graphics Suite 2017 (www.coreldraw.com).

## Acknowledgements

We gratefully thank Winfried Junke for the technical support.

## Declaration

### Availability of data and materials

All relevant data can be found within the article and its supplementary information (see Additional File 2).

### Ethics approval and consent to participate

The present study was carried out in accordance with the European Communities Council Directive of 2010 (2010/63/ EU) for care of laboratory animals and approved by a local ethics committee.

### Authors’ contributions

P.J., T.F., and S.H. conceptualized the experiments. P.J. and T.F. did the investigation and formal analysis. P.J. prepared the figures and wrote the original draft with review and editing from S.H.. All authors reviewed the manuscript.

## Additional Files

<u>Additional File 1.pdf;</u> The file shows our camera calibration performed in Matlab (Matlab 2022b; The MathWorks Inc., USA) and the respective matrices. Additionally, we plotted the shoaling parameters of each group for direct comparison and show results of the statistical tests.

<u>Additional File 2_Raw_Data.xlsx;</u> The spreadsheet contains all data points necessary to interpret or reanalyze our results. It includes individual tracking profiles and gives the data points used for statistical analysis.

<u>Additional File 3.mp4;</u> The video shows shoaling behavior of a group of flashlight fish *Anomalops katoptron* in the laboratory. Both camera perspectives, that were necessary for our analysis, are included.

