## Supplemental File 1 for "Correlation between bioluminescent blinks and swimming behavior in the splitfin flashlight fish *Anomalops katoptron*"

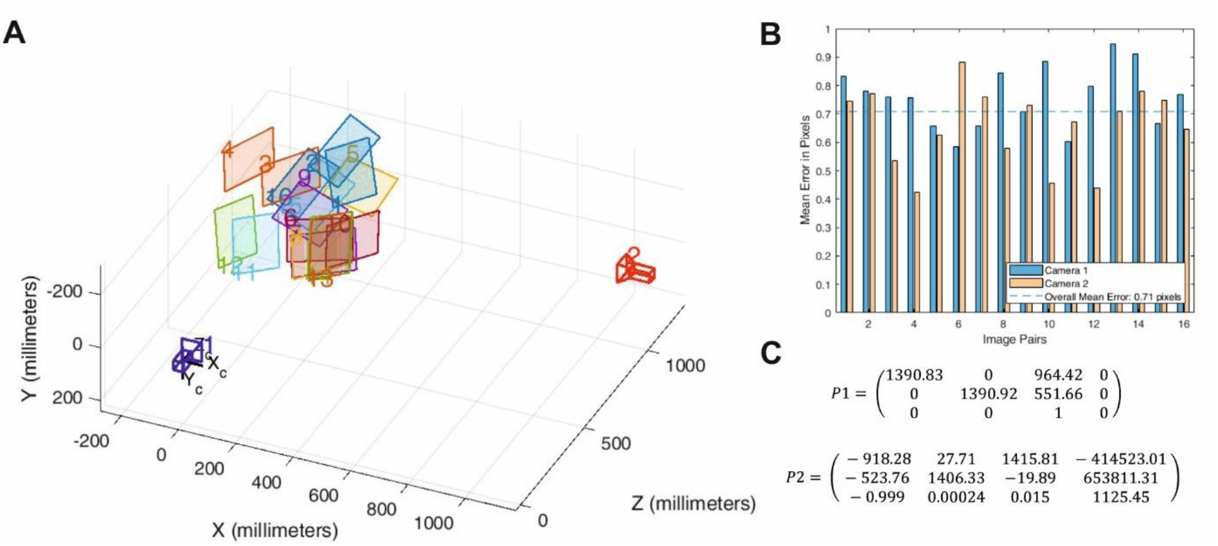


**Figure S1: Camera Parameters.** Orientation of cameras and schematic representations of the locations of calibration boards (A) and mean reprojection errors of calibration images (B) both generated with the Stereo Camera Calibration Toolbox (Matlab 2022b; The MathWorks Inc., USA). The calculated projection matrices (C) with Camera 1 (purple camera, a) being the center of the reconstructed scene.

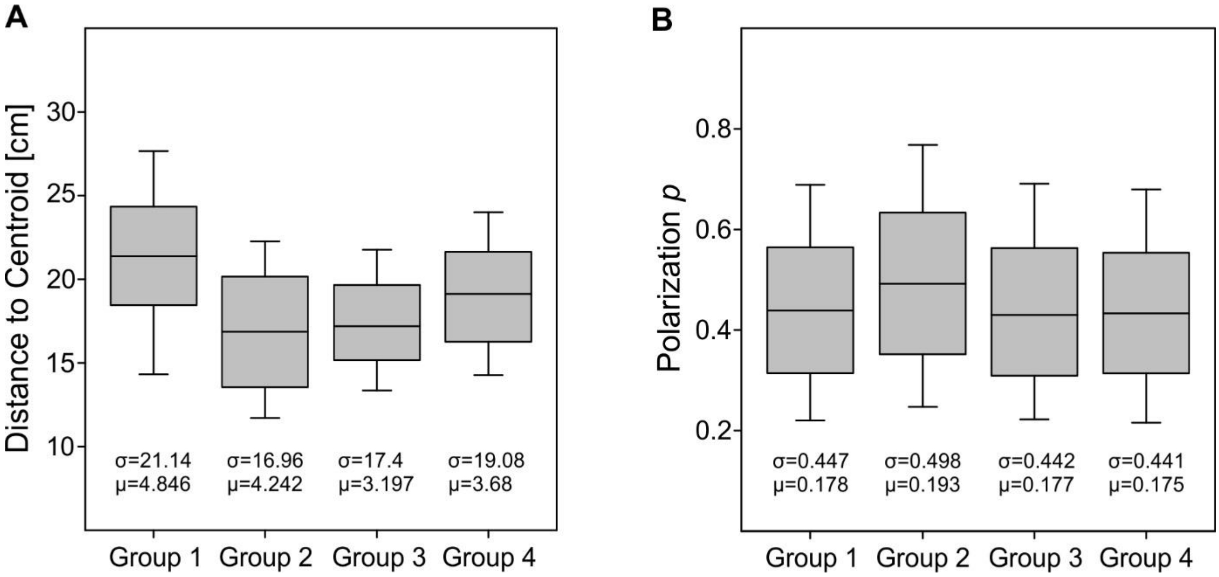


**Figure S2: Parameters for the description of group behaviors in shoals of *A. katoptron*.** In our study, four groups each consisting of five individuals were tested. To describe the shoaling behavior, we determined the mean distance to the group's centroid (A) and the polarization (B), a measure of alignment the groups individuals. Values *p* = 1 indicate the maximum alignment of all members of the group. Below the boxplot, mean (*σ*) and standard deviation (*μ*) are shown.

Figures were created with SigmaPlot 12.0 and processed with CorelDraw Graphics Suite 2017.

**Table S1: Statistical analysis of turning angle.** Shown are the results for the two-way rm-ANOVA with Holm-Sidak post-hoc analysis (see Figure 3E). We analyzed turning angle at the six timesteps (three before and after light organ transition). We used timestep and type of transition (either exposed to occluded light organs or vice versa) as factors. Statistical significance was calculated in SigmaPlot 12.0.

**Normality Test (Shapiro-Wilk)** Passed (P = 0.415)
**Equal Variance Test:** Passed (P = 0.213)

| **Source of Variation** | **DF** | **SS** | **MS** | **F** | **P** |
|---|---|---|---|---|---|
| Individual | 19 | 6007.952 | 316.208 | | |
| Switch | 1 | 1491.361 | 1491.361 | 184.70 | <0.001 |
| Switch x Individual | 19 | 153.773 | 8.093 | | |
| Frame | 5 | 1584.837 | 316.967 | 50.834 | <0.001 |
| Frame x Individual | 95 | 592.361 | 6.235 | | |
| Switch x Frame | 5 | 3528.925 | 705.785 | 63.613 | <0.001 |
| Residual | 95 | 1054.027 | 11.095 | | |
| Total | 239 | 14413.236 | 60.306 | | |

The effect of different levels of Switch depends on what level of Frame is present. There is a statistically significant interaction between Switch and Frame. (P = <0.001)

All Pairwise Multiple Comparison Procedures (Holm-Sidak method): Overall significance level = 0.05

Comparisons for factor: **Switch within 1**

| **Comparison** | **Diff of Means** | **t** | **P** | **P<0.05** |
|---|---|---|---|---|
| 2 vs. 1 | 6.815 | 6.621 | <0.001 | Yes |

Comparisons for factor: **Switch within 2**

| **Comparison** | **Diff of Means** | **t** | **P** | **P<0.05** |
|---|---|---|---|---|
| 2 vs. 1 | 1.624 | 1.578 | 0.117 | No |

Comparisons for factor: **Switch within 3**

| **Comparison** | **Diff of Means** | **t** | **P** | **P<0.05** |
|---|---|---|---|---|
| 1 vs. 2 | 12.540 | 12.183 | <0.001 | Yes |

Comparisons for factor: **Switch within 4**

| **Comparison** | **Diff of Means** | **t** | **P** | **P<0.05** |
|---|---|---|---|---|
| 1 vs. 2 | 14.867 | 14.444 | <0.001 | Yes |

Comparisons for factor: **Switch within 5**

| **Comparison** | **Diff of Means** | **t** | **P** | **P<0.05** |
|---|---|---|---|---|
| 1 vs. 2 | 8.189 | 7.956 | <0.001 | Yes |

Comparisons for factor: **Switch within 6**

| **Comparison** | **Diff of Means** | **t** | **P** | **P<0.05** |
|---|---|---|---|---|
| 1 vs. 2 | 2.757 | 2.678 | 0.009 | Yes |

**Table S2: Statistical analysis of swimming speed.** Shown are the results for the two-way rm-ANOVA with Holm-Sidak post-hoc analysis (see Figure 3F). We analyzed swimming speeds at the six timesteps (three before and after light organ transition). We used timestep and type of transition (either exposed to occluded light organs or vice versa) as factors. Statistical significance was calculated in SigmaPlot 12.0.

**Normality Test (Shapiro-Wilk)** Passed (P = 0.105)
**Equal Variance Test:** Passed (P = 0.356)

| Source of Variation | DF | SS | MS | F | P |
|---|---|---|---|---|---|
| Individual | 19 | 0.628 | 0.033 | | |
| Switch | 1 | 0.053 | 0.053 | 143.075 | <0.001 |
| Switch x Individual | 19 | 0.007 | 0.0003 | | |
| Frame | 5 | 0.047 | 0.0094 | 41.917 | <0.001 |
| Frame x Individual | 95 | 0.021 | 0.0002 | | |
| Switch x Frame | 5 | 0.182 | 0.0364 | 38.719 | <0.001 |
| Residual | 95 | 0.089 | 0.0009 | | |
| Total | 239 | 1.029 | 0.0043 | | |

The effect of different levels of Switch depends on what level of Frame is present. There is a statistically significant interaction between Switch and Frame. (P = <0.001)

All Pairwise Multiple Comparison Procedures (Holm-Sidak method): Overall significance level = 0.05

Comparisons for factor: **Switch within 1**

| Comparison | Diff of Means | t | P | P<0.05 |
|---|---|---|---|---|
| 2 vs. 1 | 0.0643 | 6.993 | <0.001 | Yes |

Comparisons for factor: **Switch within 2**

| Comparison | Diff of Means | t | P | P<0.05 |
|---|---|---|---|---|
| 2 vs. 1 | 0.0862 | 9.372 | <0.001 | Yes |

Comparisons for factor: **Switch within 3**

| Comparison | Diff of Means | t | P | P<0.05 |
|---|---|---|---|---|
| 2 vs. 1 | 0.0885 | 9.621 | <0.001 | Yes |

Comparisons for factor: **Switch within 4**

| Comparison | Diff of Means | t | P | P<0.05 |
|---|---|---|---|---|
| 2 vs. 1 | 0.0238 | 2.588 | 0.011 | Yes |

Comparisons for factor: **Switch within 5**

| Comparison | Diff of Means | t | P | P<0.05 |
|---|---|---|---|---|
| 1 vs. 2 | 0.0377 | 4.103 | <0.001 | Yes |

Comparisons for factor: **Switch within 6**

| Comparison | Diff of Means | t | P | P<0.05 |
|---|---|---|---|---|
| 1 vs. 2 | 0.0463 | 5.038 | <0.001 | Yes |

**Additional File 2**

Additional File 2_Raw_Data.xlsx

The spreadsheet contains all data points necessary to interpret or reanalyze our results. It includes individual tracking profiles and gives the data points used for statistical analysis.

**Additional File 3**

Additional File 3.mp4

The video shows shoaling behavior of a group of flashlight fish *Anomalops katoptron* in the laboratory. Both camera perspectives, that were necessary for our analysis, are included.
